# Unexplored Yeast diversity in Seed Microbiota

**DOI:** 10.1101/2024.11.27.625647

**Authors:** Muriel Marchi, Anaïs Bosc-Bierne, Laurine Labourgade, Thomas Lerenard, Josiane Le Corff, Sophie Aligon, Aurélia Rolland, Marie Simonin, Coralie Marais, Martial Briand, Viviane Cordovez, Linda Gouka, Thomas Guillemette, Philippe Simoneau, Natalia Guschinskaya

## Abstract

Yeasts are known to be fantastic biotechnological resources for medical, food, and industrial applications, but their potential remains untapped in agriculture, especially for plant biostimulation and biocontrol. In particular, yeasts have been reported as part of the core microbiome of seeds using next generation sequencing methods, but their diversity and functional roles remain largely undescribed. Focusing on yeasts and excluding filamentous fungi, this study aimed to characterize the diversity of seed-associated yeasts across nine plant species (crops and non-cultivated species) using culturomics and microscopy. Comparison with available metabarcoding data was performed to assess the representativeness of the strain collection in seed samples. Our results show that seed-associated yeasts largely belong to Basidiomycota phylum and more particularly to the Tremellomycetes class. This yeast collection covers 15 genera (2 of Ascomycota and 13 of Basidiomycota). Out of the 229 isolates described, the most frequently isolated yeasts were *Holtermanniella*, *Vishniacozyma*, *Filobasidium, Naganishia* and *Sporobolomyces.* The yeasts from these dominant genera were isolated from multiple plant species (4 to 8), except for *Naganishia* which only originated from *Solanum lycopersicum L*. These results are also consistent with the fact that these dominant taxa were recently identified as members of the core seed microbiome, indicating their high prevalence and abundance across diverse plant hosts and environments. Compared to previous plant yeast diversity surveys, the members from Ascomycota yeasts are less frequent in seeds and only represented here by the *Aureobasidium* and *Taphrina* genera. Altogether, these results suggest that yeasts are generally well-adapted to the aboveground habitats of plants, but seeds represent a specific habitat that diverse Basidiomycota yeasts can colonize.

**Take away message:**

- 229 yeasts isolated from seeds and seedlings of diverse plant species
- Most isolates are Basidiomycota yeasts, especially of the Tremellomycetes class
- The most frequently isolated yeasts belong to *Holtermanniella*, *Vishniacozyma*, *Filobasidium* and *Sporobolomyces* genera
- The collection is representative of taxa found in seed microbiota of multiple plant species, including core members

## Introduction

There is increasing awareness of the role of plant microbiota in plant fitness and health (Compant et al., 2016; Trivedi et al., 2020). Many examples of the beneficial effects of plant-microbiome interactions originate from studies on the rhizosphere and phyllosphere, demonstrating higher nutrient acquisition, abiotic stress tolerance or disease resistance (de la Fuente Cantó et al., 2020; Vannier et al., 2019). More recently, research has been conducted on the diversity and role of seed microbiota, which has so far been neglected despite its role as a primary inoculum of plants (Abdelfattah et al., 2023; Bergmann et al., 2023; Shade et al., 2017).

Seeds are crucial for crop settlement and food production. However, only a limited number of studies have investigated the role of seed-associated microorganisms in plant germination, emergence, and tolerance to soil-borne or seed-transmitted pathogens (Figueiredo dos Santos et al., 2021; Lamichhane et al., 2018; Matsumoto et al., 2021; Rochefort et al., 2019; Simonin et al., 2023). Moreover, the available studies have focused primarily on the seed-associated bacterial community, ignoring the highly diversified fungal component. Indeed, a specificity of the seed habitat is to host similar levels of fungal and bacterial diversity, whereas other plant or soil habitats are generally dominated by bacterial taxa (Johnston-Monje et al., 2021; Simonin et al., 2022). A recent meta-analysis of seed microbiota across 50 plant species identified over 2,000 seed fungal taxa, as determined by the ITS1 region. This includes 16 core taxa that are present in more than 20 plant species, as well as flexible taxa that are primarily specific to certain plant and seed production areas (Simonin et al., 2022). These core taxa were extremely abundant in most samples and were mainly affiliated with the filamentous fungi genera, *Cladosporium* and *Alternaria*. However, an interesting observation was that six seed core taxa were yeasts, affiliated with the genera *Vishniacozyma*, *Filobasidium*, *Sporobolomyces*, and *Aureobasidium*. DNA of isolates from these genera are commonly detected in various environments and biomes as soil, root, shoot, air, litter from croplands, forest, deserts (Vetrovsky et al., 2020).

Yeasts are widely recognized as valuable biotechnological resources in medical, food, and industrial applications. However, their potential in agriculture, particularly for plant biostimulation and biocontrol, remains underexplored, except for their use in post-harvest management (Ezzouggari et al., 2024; Freimoser et al., 2019; He et al., 2024).

Despite their frequent detections as core or hub taxa in various plant habitats using next generation sequencing methods (Agler et al., 2016; Almario et al., 2022; Noel et al., 2022; Simonin et al., 2022) only few studies are available on plant yeasts. They have been mainly conducted in the phyllosphere, nectar, and fruit compartments (Gouka et al., 2022; Jacquemin et al., 2020) indicating adaptation to aboveground plant habitats. Still, characterization of the diversity and functions of yeast colonizing seeds has received even less attention, with few yeast isolates, genomes, or functional studies available (Bhuyan et al., 2023; Nelson et al., 2018).

This study aimed to fill one of these knowledge gaps by characterizing the diversity of seed-associated yeasts across nine plant species (crops and non-cultivated species) using culturomics and microscopy. The following questions were addressed: What is the taxonomic diversity of the yeast isolated from seeds? Which yeast taxa are shared or specific to a given plant species? How is our yeast collection representative of yeast diversity detected by metabarcoding analyses (i.e., core and flexible taxa)?

Fungal isolation was performed on seeds from three wild Brassicaceae species (*Capsella bursa-pastoris*, *Draba verna* and *Cardamine hirsuta*), and on seeds and seedlings (grown *in vitro*) from five crops (*Phaseolus vulgaris*, *Raphanus sativus*, *Brassica napus* L., *Solanum lycopersicum* L. and *Triticum aestivum*). Taxonomic and morphological identification of yeasts was conducted using ITS/D1-D2 sequencing, combined with macroscopic and microscopic observations.

We then determined if the yeast taxa isolated in our collection were commonly found in seed samples using the existing Seed Microbiota Database based on ITS1 metabarcoding. This analysis allowed us to assess the retrieval of core, abundant, and rare yeast taxa using culturomics.

## Material and Methods

### Seed material

Seeds from five crops and three wild *Brassicaceae* were used for yeast isolation. The selection of plant species in this study covers major crops from four main cultivated botanical families (Poaceae, Fabaceae, Brassicaceae, Solanaceae) and from wild species present in contrasted land-uses (urban to field). This collection of seed material came from distinct research projects. Crops seeds (*Phaseolus vulgaris*, *Brassica napus* L., *Solanum lycopersicum* L. and *Triticum aestivum*) were produced in different conditions (field, nursery, mini cage, greenhouse) from various French’s areas (IGEPP, Le Rheu, Bretagne (48.13916, −1.799776); FNAMS, Loire Authion, Anjou (47.470377, −0.394989); FNAMS, Condom, Gascogne (43.956065, 0.3921756470); GAFL, Avignon, Provence; GDEC, Clermont Ferrand, Auvergne). Wild *Brassicaceae* (*Capsella bursa-pastoris*, *Draba verna*, *Cardamine hirsuta*) were harvested in three different ecotypes from Pays de la Loire (field, peri-urban, urban: 47°25’38.0”N 0°31’48.3”W, 47°28’51.0”N 0°36’30.7”W, 47°25’18.3”N 0°34’42.4”W, 47°25’34.9”N 0°33’54.0”W). The *Raphanus sativus* seed lot was a commercial lot, no information was available. More details about the plant material are listed in Table S1.

### Isolation procedure

Taking advantage of our various laboratory strain collections, the yeast isolates presented in this paper originate from three main isolation campaigns, each carried out within separate scientific projects with distinct objectives. This context explains the differences in both isolation methods and the types of materials used (seeds and seedlings). The aim of this paper is not to compare the isolation methods, but rather to explore the diversity of yeasts within the seed microbiota across our fungal collection.

Yeasts were therefore isolated from two types of biological material (seeds and seedlings), using two methods: i) placing seeds directly on agar plates or ii) using macerates of seeds and seedlings germinated in sterile conditions, as described below.

#### Yeast isolation methods

##### Wild Brassicaceae

One hundred seeds from each of the wild *Brassicaceae* plants were placed in 20 Petri dishes (90-mm diameter). Ten Petri dishes contained a Potato Dextrose Agar (PDA) (39 g/L, BIOKAR Diagnostics) including streptomycin (500 mg/L, Sigma-Aldrich) and 10 dishes contained a semi-selective medium adapted from Dhingra and Sinclair (1985) with PDA (39 g/L) including 150 mg/L of Rose Bengal and 250 mg/L of chloramphenicol (Sigma-Aldrich). Plates were incubated at 22°C in the dark and monitored daily to collect yeasts (Lerenard et al., 2024). For these Brassicaceae species only seeds, not seedlings, were used for isolation, due to the limited amount of seed available.

##### Crops

Seeds from crop species and seedlings germinated in sterile conditions (as described by Barret et al., 2015) were soaked in a solution of Phosphate Buffered Saline including 0.05% of Tween 20 (Fisher Scientific, 12673967) in order to collect both seed epiphytes and endophytes microorganisms following International Seed Testing Association (ISTA) guidelines. The seedlings germinated *in vitro* enable us to identify and isolate seed-borne yeasts that have the capacity to transmit to developing plants, surviving the strong bottleneck imposed by germination and seedling emergence (Barret et al., 2015). Three biological repetitions were used for *Solanum lycopersicum L.* (including 1g, approx. 360 seeds and 6 seedlings), *Triticum aestivum* (including 4g, approx. 100 seeds and 6 seedlings), *Raphanus sativus* (including 4 g, approx. 1000 seeds) and *Brassica napus L.* (including 2 g, approx. 400 seeds and 15 seedlings) and five biological repetitions for *Phaseolus vulgaris* (including 4 to 18 g, approx. 30 seeds and 30 seedlings).

Maceration volumes for seeds were 4, 8, 8 and 4 mL for *Solanum lycopersicum L.*, *Triticum aestivum*, *Raphanus sativus* and *Brassica napus L.* respectively. For *Phaseolus vulgaris*, given the large size variation of the seeds, the maceration volume was 2 mL/g of seeds (ranging from 7 to 36 mL). Seed maceration was conducted under agitation at 6°C for 2h30 for *Raphanus sativus* and *Brassica napus L.,* and 16h overnight for *Solanum lycopersicum L.*, *Triticum aestivum* and *Phaseolus vulgaris.* For *Solanum lycopersicum L*. and *Triticum aestivum*, after maceration, the seeds were placed in a stomacher machine for 3 minutes. For these plant species, seedlings were crushed in 2 mL sterile water to obtain a homogeneous solution used for the filtration and isolation process.

For yeast strain isolation, the seed and seedling macerates obtained were filtered using filtration units with mesh woven filters 31 µm (SEFAR NITEX 03-31/24). The supernatant containing the microorganisms that went through the filters (mainly bacteria) was discarded to isolate only larger organisms present on the filters (yeast and filamentous fungi). Thus, we recognize the potential for selective bias introduced by this filtration step. But it proved effective for our objective of isolating diverse fungal taxa from plant material under reduced bacterial growth pressure.

The filter was washed in 5 ml of sterilized osmotic water. The suspension was diluted in liquid malt 1.5% with tetracycline 10 mg/L (Sigma-Aldrich) and streptomycin 50 mg/L (Sigma-Aldrich) to prevent bacterial growth and dilutions (1/10, 1/50, 1/100, 1/200 and 1/500) were done in 96-well plates. Plates were then incubated at 23°C for one week. The following steps of the isolation procedure focused on microplates presenting microbial growth in half or less than half of the wells to maximize the chance to have pure culture in each well (Zhang et al., 2021). When a well-presented growth, the microbial suspension was collected using a micropipette and deposited on a 1.5% malt agar plate. Plates were incubated at 23°C until the observation of fungal development. Non-filamentous isolates were selected, and yeast’s shapes were checked using a bright-field microscope. Yeast isolates were purified, and strains were stored at −80°C in glycerol 30%.

### PCR-based method (ITS and D1D2) for yeast identification

Yeast identification was carried out based on the sequence of the D1/D2 region of the large 26S rDNA subunit using primers LROR (5’-ACCCGCTGAACTTAAGC-3’) and LR5 (5’-TCCTGAGGGAAACTTCG-3’) (Vilgalys & Hester, 1990) and the internal transcribed spacer (ITS) region using primers NS7 (5’-GAGGCAATAACAGGTCTGTGATGC-3’) and ITS4 (5’-TCCTCCGCTTATTGATATG-3’) (White et al., 1990). Gene sequences were amplified by colony PCR as described by Mirhendi (Mirhendi et al., 2007). We used GoTaq® G2 Hot Start Polymerase kit from Promega Corporation in concordance with manufacturer’s instructions for PCR amplification and Sanger sequencing using Eurofins Genomics facilities.

### Sequence treatment and tree inference

The details of the reference sequences used in this study, including the accession numbers and their literature origins, are listed in Table S2.

All sequence alignments were produced with the server version of MAFFT (Katoh et al., 2019), with a gap open penalty of 5 and a gap extension penalty in the range of 0.05 to 1. After trimming the ITS and D1/D2 alignments, the concatenated sequences were used for phylogenetic analysis. The model used for the tree inference was a general time reversible model with unequal rates and unequal base frequency (GTR+F+I+G4). The maximum likelihood phylogenetic tree was built using IQ-TREE web server (Trifinopoulos et al., 2016) and branch support with the ultrafast bootstrap (Hoang et al., 2018). The figure was obtained using Interactive Tree Of Life (iTOL)v6 (Letunic & Bork, 2021).

### Yeast Morphological Description

Descriptions of agar-grown yeast colonies were done on Yeast Malt extract Agar (YMA) plate including texture, colour, surface, elevation and margin as described by (Kurtzman et al., 2011). Culture colour assessments were subjectively coded based on observations from 10-day-old cultures on YMA at 22°C in the dark.

Microscopic images for each yeast strain were obtained using a transmission microscope with Differential Interference Contrast (DIC) optics (Axio Imager Z2, Carl Zeiss, Oberkochen, Germany) and ZenBlue software at x100 magnification (ROI 1024x). For sample preparation for each isolate, a single colony was picked with a sterile toothpick and transferred into a tube containing 50μL of sterile water. A 20μL drop of suspension was then placed between a microscope slide and coverslip for observation. Figure 2 presents macroscopic and microscopic photographs of a representative isolate of each genus.

### Prevalence and abundance of yeast isolates in seed microbiota

To determine if the yeast taxa isolated in our collection were commonly found in seed samples, we took advantage of the existing Seed Microbiota Database (Simonin et al., 2022). This database compiles metabarcoding data from multiple seed microbiota studies, including 17 studies that characterized fungal diversity using ITS1 marker, generating data from 1037 seed samples and 34 plant species. We sequenced the NS7–ITS4 region of the fungal isolates in our collection using Sanger sequencing. Only high-quality sequences were used for the following analysis (n=219) excluding sequences with non-identified nucleotide. From these high-quality sequences, we generated *in silico* Amplicon Sequence Variants (ASVs) by trimming them to match exactly with the ITS1f/ITS2 region (229 bp) used in the metabarcoding study by Simonin et al. (2022). We obtained the final ITS1 Amplicon Sequencing Variant (ASV) corresponding to each of our isolates. This allowed for a direct comparison between our cultured isolates and the ASVs reported in this large-scale meta-analysis. This comparison enabled us to evaluate the representativeness of our collection within the broader seed mycobiota landscape.

Because of the short ITS1 fragment considered (genus-level resolution), several isolates possessed the same ASV and for all isolates we obtained a total of 36 unique ASVs. We then determined for each isolate ASV: the prevalence (number of samples detected), the relative abundance, and the number of plant species in which they were detected in the Seed Microbiota Database. We also compared them to the list of “core” fungal taxa identified in this previous study. Here, ASVs were identified as core taxa if they were detected in ≥ 20 plant species.

The script for the analysis and figures generated in R is available here: https://github.com/marie-simonin/Yeast_Diversity_Seed_Microbiota.

## Results

### Overview of the Fungal Collection

As explained in the “Materials and Methods” section, the collection was obtained from separate research projects with different objectives and isolation procedures. A total of 539 fungi were obtained from wild Brassicaceae seeds, including 49 yeasts (approximately 9% of the total). From crops seeds and seedlings, 1020 isolates were obtained, including 242 yeasts (191 from seeds and 51 from seedlings, or approximately 24% of the isolates). The isolation campaigns were carried out on different seed lots and with two different methods. Therefore, we will not compare these results obtained with both methods. However, our observations suggest that the isolation method using macerates could increase the recovery rate of yeast isolates. The following analysis focuses only on the yeasts in the collection for which molecular typing was successful (n = 229).

### Isolation of a large diversity of yeasts from seeds and seedlings

A total of 229 yeasts were identified. Phylogenetic tree inference (Figure 1) gathers a total of 362 strains including reference strains from several studies listed in the table S2.

**Figure 1:**
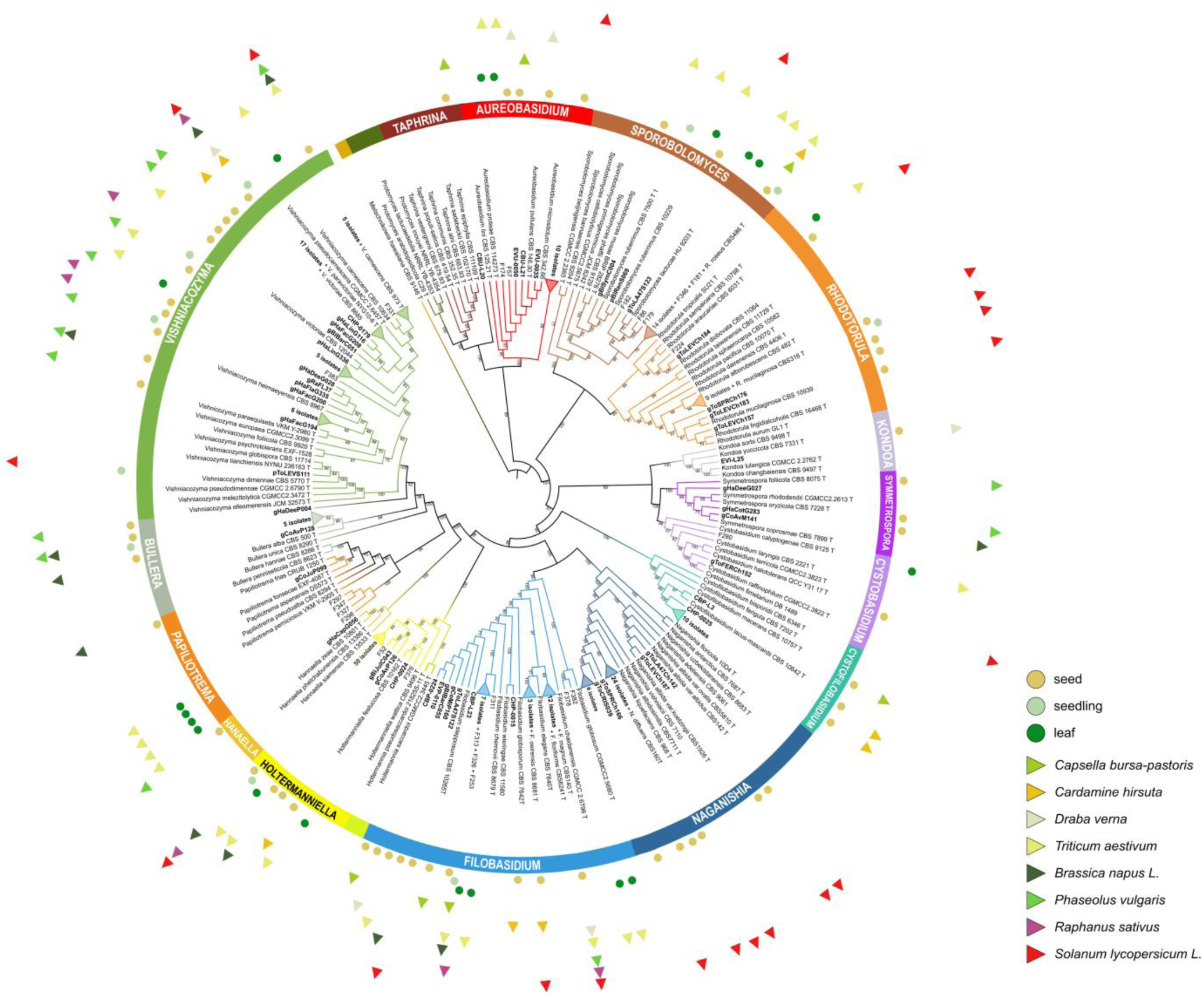
Circular phylogenetic tree of the 229 strains from seed and seedlings yeast collection. Strains isolated on seed (full brown circle) of *Capsella bursa-pastoris* (light green triangle), *Draba verna* (light grey triangle) and *Cardamine hirsuta* (orange triangle), and on seeds and seedlings (full light green circle) from five crops *Phaseolus vulgaris* (green triangle), *Raphanus sativus* (purple triangle), *Brassica napus L.* (black triangle), *Solanum lycopersicum L.* (red triangle), and *Triticum aestivum* (yellow triangle). *Metchnikowia* stains were used as an outgroup. The tree includes also some strains from Gouka et al. (2022), isolated from *Triticum aestivum* phyllosphere (yellow triangle for *Triticum aestivum* and dark green circle for the leaf). Phylogeny is based on the alignment and the concatenation of ITS and D1/D2 domain partial sequences. The alignments were produced with the server version of MAFFT (Katoh et al., 2019), and the maximum likelihood phylogenetic analysis using IQ-TREE web server (Trifinopoulos et al., 2016) and branch support with the ultrafast bootstrap (Hoang et al., 2018). The tree figure was obtained using Interactive Tree Of Life (iTOL)v6 (Letunic & Bork, 2021).

All the yeasts belong to the fungi kingdom to the phyla Ascomycota and Basidiomycota. Our yeast collection includes 2 genera from Ascomycota phylum (with n=14 strains) and 13 genera from Basidiomycota phylum (with n=215) (**Figure 1**, **Table 1 and Table S1**).

**Table 1:**
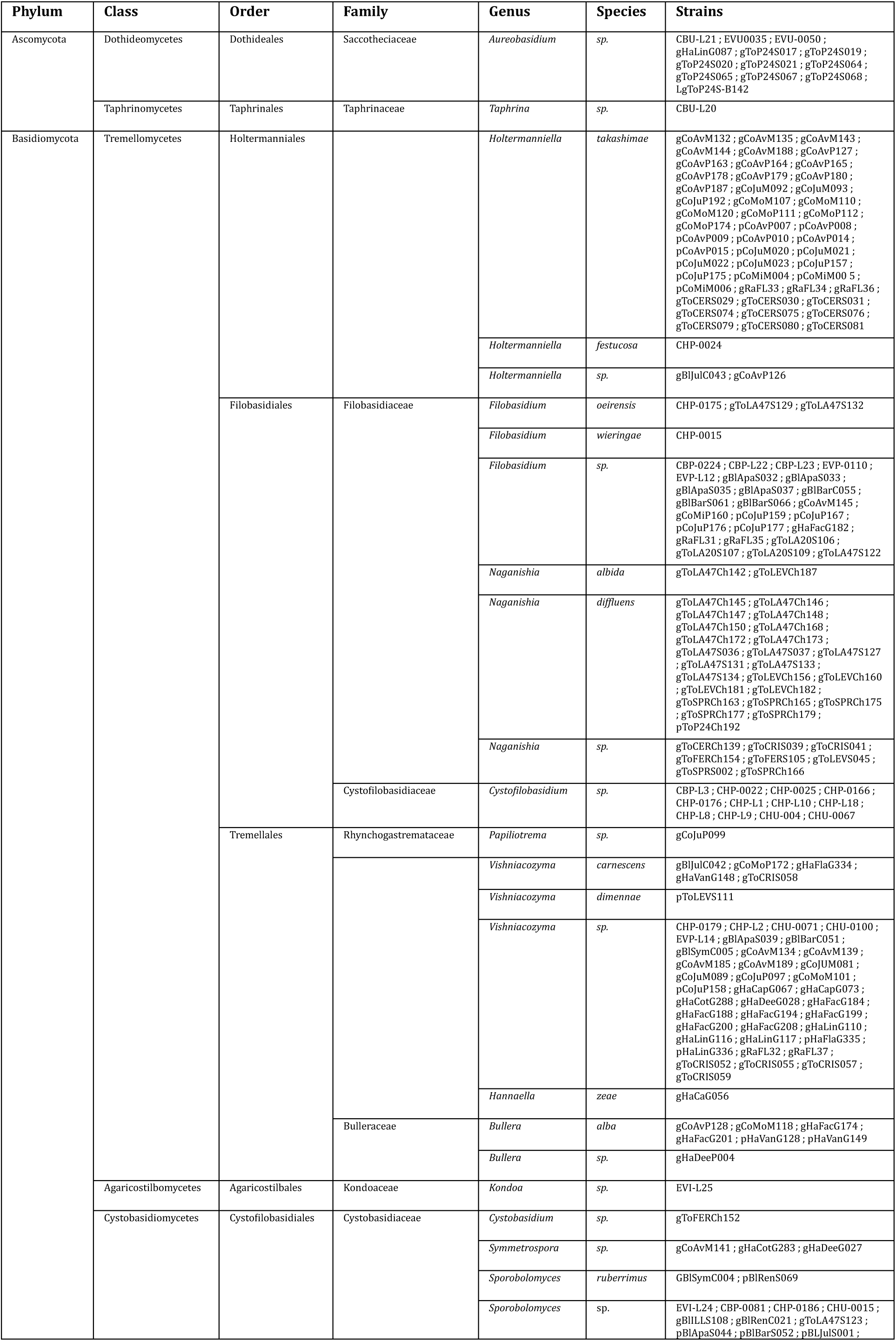
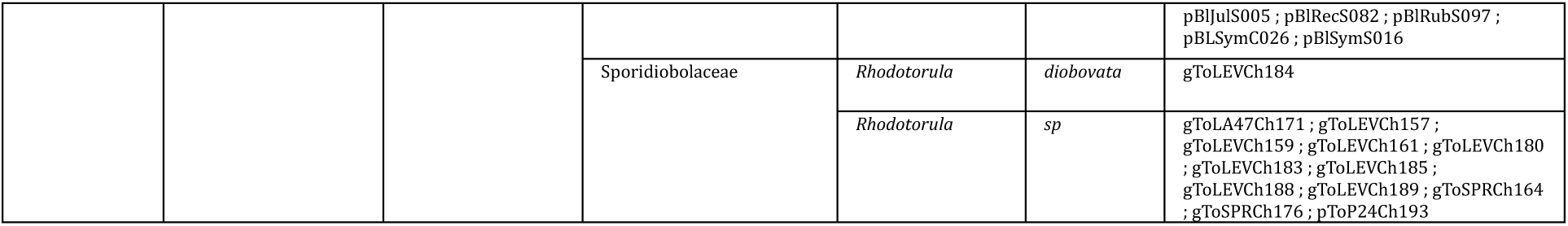
List of yeast taxa and species identified in the seed collection.

Four genera were highly represented in our collection: *Holtermanniella* (52 isolates), *Vishniacozyma* (44 isolates), *Naganishia* (34 isolates) and *Filobasidium* (29 isolates). The other genera are less represented: *Sporobolomyces* (17 isolates), *Cystofilobasidium* (12 isolates), *Rhodotorula* (13 isolates), *Bullera* (7 isolates), *Aureobasidium* (13 isolates), *Symmetrospora* (3 isolates), *Cystobasidium* (1 isolate), *Hannaella* (1 isolate), *Kondoa* (1 isolate), *Papiliotrema* (1 isolate) and *Taphrina* (1 isolate).

Details about isolate’s affiliation are synthetized in Table 1. All the strains were identified at the genus level. Around 42% of them were affiliated at species level (n=96) and for 133 isolates the identification is reliable only at the genus level. The main reason for this result is due to the lack of similar sequences in GenBank (for example, CBU-L20) or the lack of resolution of the selected genetic markers (for example *Filobasidium floriforme* or *F. magnum*) (Vu et al., 2016).

### Macroscopical and microscopical strains description

Figure 2 provides a description of representative isolates (one from each genus) from our yeast collection at both macroscopic and microscopic scales. This description is complemented by detailed colony morphology presented in Table S1.

**Figure 2:**
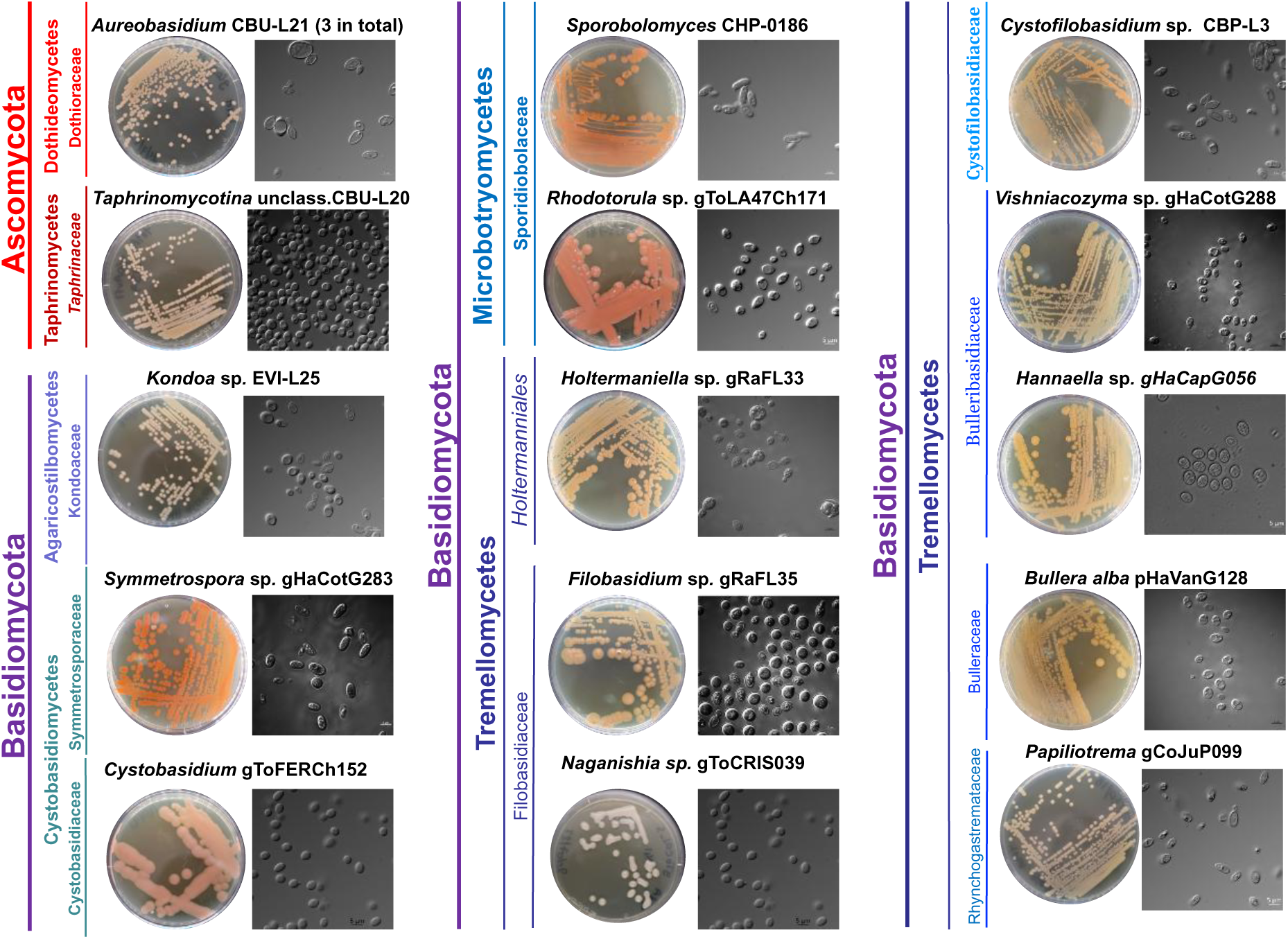
Phenotypic characterization of the selected yeast isolates on a macroscopical and microscopical scale. One isolate is presented for each genus. Strains are classified by phylum, class, family, and genus. The pictures of agar-grown yeast colonies on Yeast Malt extract Agar (YMA) were taken after 10 days of incubation at 22°C in the dark. Microscopic images for each yeast type strain were obtained using a transmission microscope with Differential Interference Contrast (DIC) optics x100, ROI 1024, and ZenBlue software (Axio Imager Z2, Carl Zeiss, Oberkochen, Germany), scale 5 µm.

Most colonies cultured on YMA exhibited a butyrous texture. Notably, some colonies displayed mucoid growth (as gToFERCh152 from *Cystobasidium* genus and gToCRIS039 from *Naganishia* genus), often linked to elevated production of extracellular polysaccharides (refer to Table S1 for examples). Additionally, rare occurrences of yellow non-diffusible carotenoid pigments were observed in the isolates from genera *Holtermanniella, Cystofilobasidium* and *Hannaella* (Figure 2 and Table S1). Interestingly, 15 isolates (from *Symmetrospora*, *Sporobolomyces*, *Rhodotorula*) had red colour, the members of these three genera are known to produce red non-diffusible pigments of carotenoid composition (Chreptowicz et al., 2019; Zhao et al., 2019) (Figure 2 and Table S1). Some isolates (as gToFERCh152) were observed to form creamy colonies that may become slimy with age.

Yeasts from our collection are relatively the same size, measuring between 6-10 μm. They are generally distinguished by their round, oval, or elongated shapes, and the nucleus and organelles of each cell can be easily seen through the cell wall and cytoplasmic membrane. It is possible to observe budding yeasts and the scars they leave after binary fission of the entities (Figure 2).

### Yeast plant’s specificity and ecotype

Most yeasts from the collection were isolated from seeds (n= 193) and not from seedlings (n=36). Isolates from seed and seedling are distributed across the entire taxonomic tree (Figure 1).

Four genera were highly represented in the collection: *Holtermanniella* (52 isolates), *Vishniacozyma* (44 isolates), *Naganishia* (34 isolates) and *Filobasidium* (29 isolates). The yeasts from these dominant genera were generally isolated from multiple plant species (4 to 8), except for *Naganishia* which only originated from *Solanum lycopersicum L.* Four other genera had between 10 and 20 isolates: *Sporobolomyces* (17), *Rhodotorula* (13), *Aureobasidium* (13), *Cystofilobasidium* (12). *Sporobolomyces* isolates came from 5 plant species, *Aureobasidium* isolates from 4 plant species while *Rhodotorula* was specific to *Solanum lycopersicum L.* and *Cystofilobasidium* to wild Brassicaceae (Figure 3). The other ten genera were represented by only a few isolates (1 to 5) generally originating from a single plant species.

**Figure 3:**
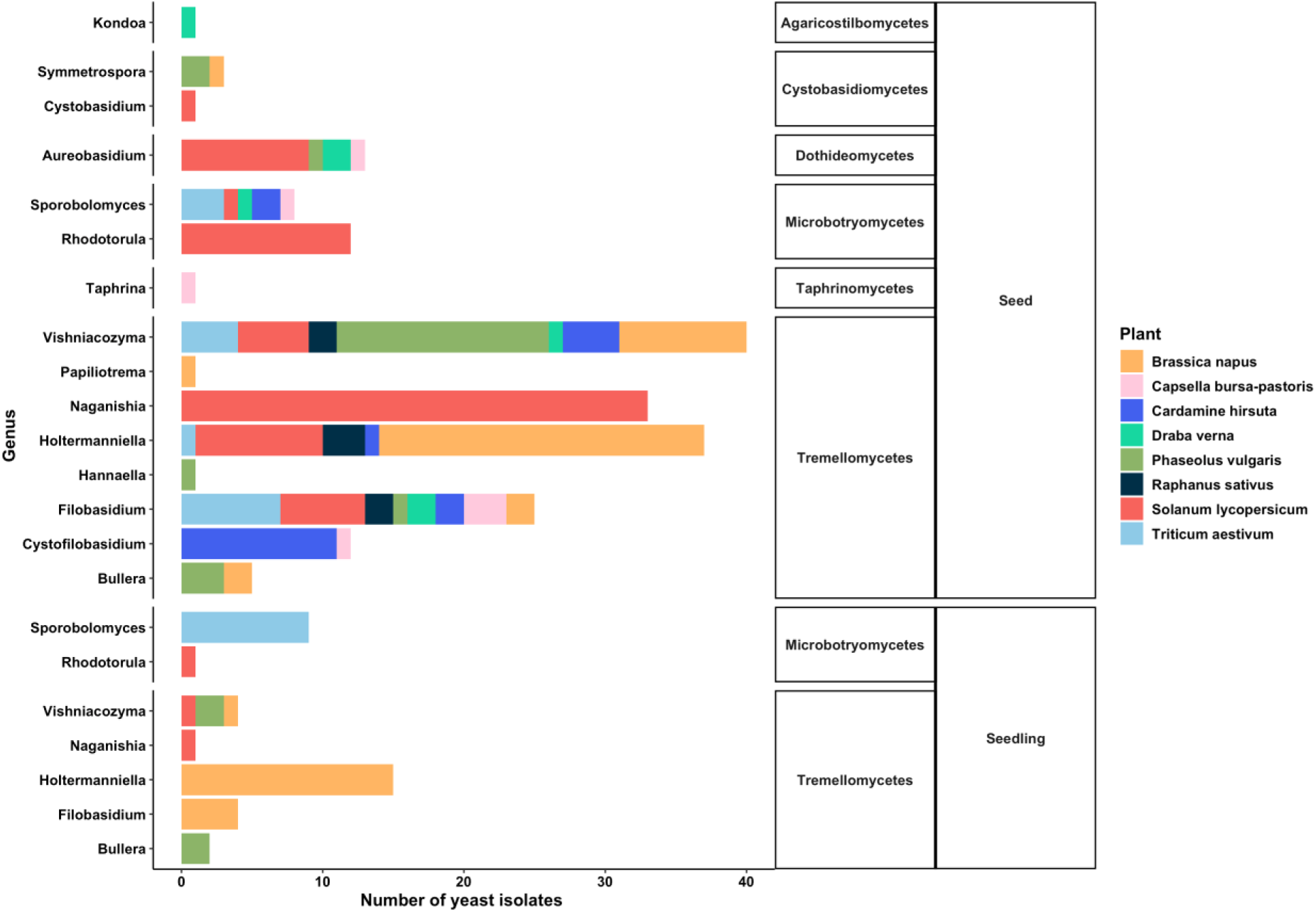
Number of yeast isolates for the 15 genera from our yeast collection, colored by their plant host and organized in function of the plant compartment of isolation and fungal class. Please note that yeast isolation on seedlings was only performed on four species (*Brassica napus*, *Phaseolus vulgaris*, *Solanum lycopersicum* and *Triticum aestivum*).

Only 7 genera were isolated from both seeds and seedlings (grown *in vitro* for 4 crops) and they were the most represented in the collection (*Holtermanniella*, *Vishniacozyma*, *Filobasidium*, *Naganishia*, *Rhodotorula*, *Sporobolomyces, Bullera*) (Figure 3).

For detailed information on the origin of each strain, its taxonomic affiliation, and sequences, see Table S1 and Table S2.

### Yeast isolates are representative of core and flexible taxa found in seed microbiota of multiple plant species

To evaluate how the yeast collection is representative of the known diversity of seed microbiota, we used a recent meta-analysis on seed microbiota diversity across 50 plant species. The analysis of prevalence and abundance enabled us to identify which isolates were present in the core taxa of the seed microbiota. Conversely, it also helped us determine which isolates could be considered specific to our seed lots and plants.

After extracting the ASVs of our isolates matching the same ITS1 fragment used in the meta-analysis (Seed Microbiota Database), we determined their prevalence, relative abundance and number of plant hosts across all seed samples available (n=1037, Figure 4).

**Figure 4:**
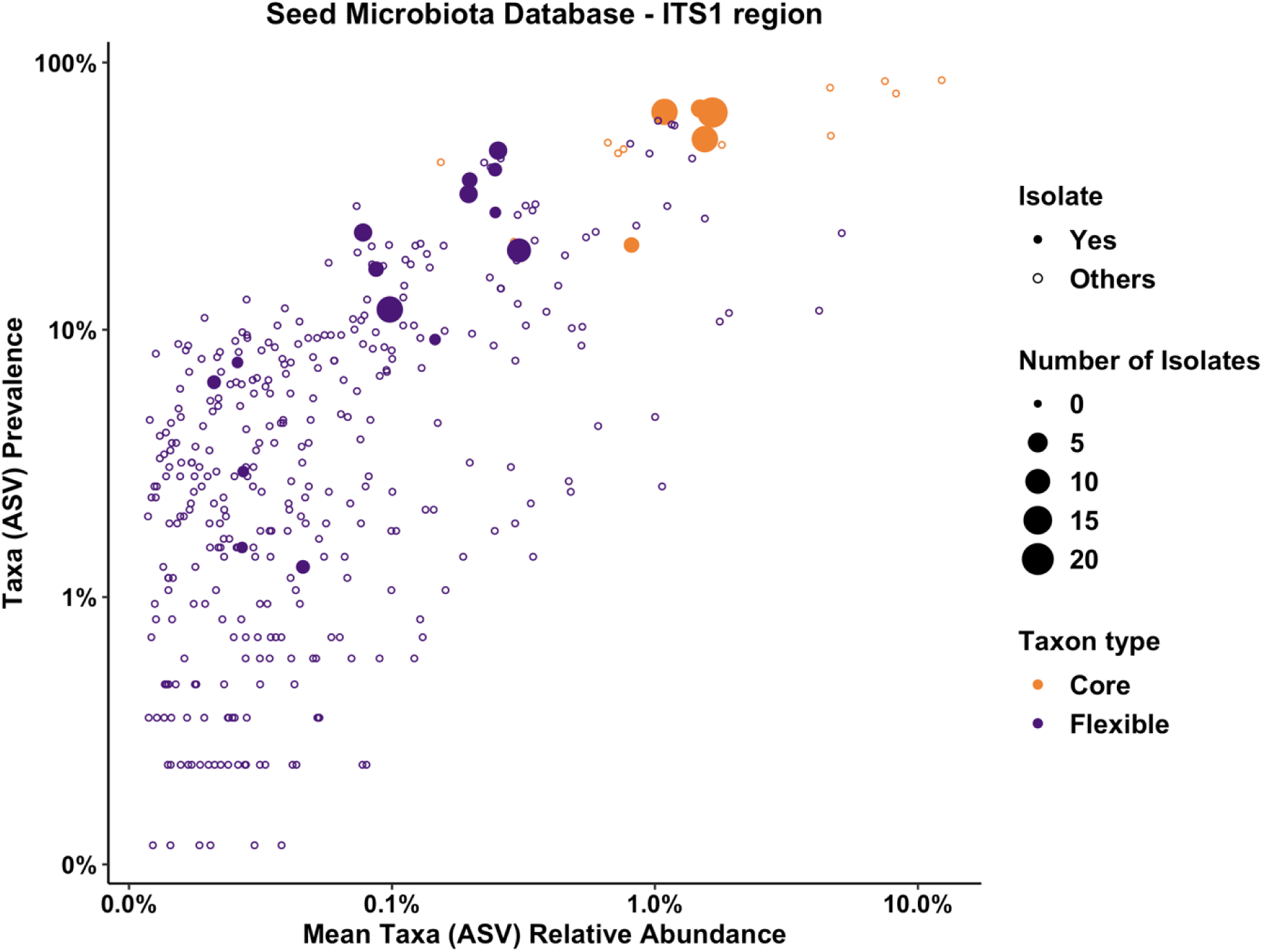
Abundance–occupancy curves of all fungal amplicon sequence variants (ASVs, including filamentous fungi and yeasts) detected in the Seed Microbiota Database (Simonin et al., 2022) using ITS1 region metabarcoding (ASVs < 100 reads were excluded). Each point represents an ASV, and the solid circles indicate ASVs for which we have yeast isolates, the points are coloured based on their characterization as a core (detected in > 20 plant species) or flexible taxon. The size of the point is proportional to the number of yeast isolates in the collection.

The yeast isolates represented 36 ASVs of which 25 were detected in the meta-analysis (list of detected and non-detected ASVs in Table 2). The isolate ASVs covered a wide range of abundance-occupancy patterns, ranging from ubiquitous (> 60% prevalence, i.e. present in > 60% of all samples studied) to rare taxa (<1% prevalence). All ASVs were detected in multiple plant species, except two, suggesting the frequent associations of these taxa with seeds originating from various hosts and environments.

**Table 2:**
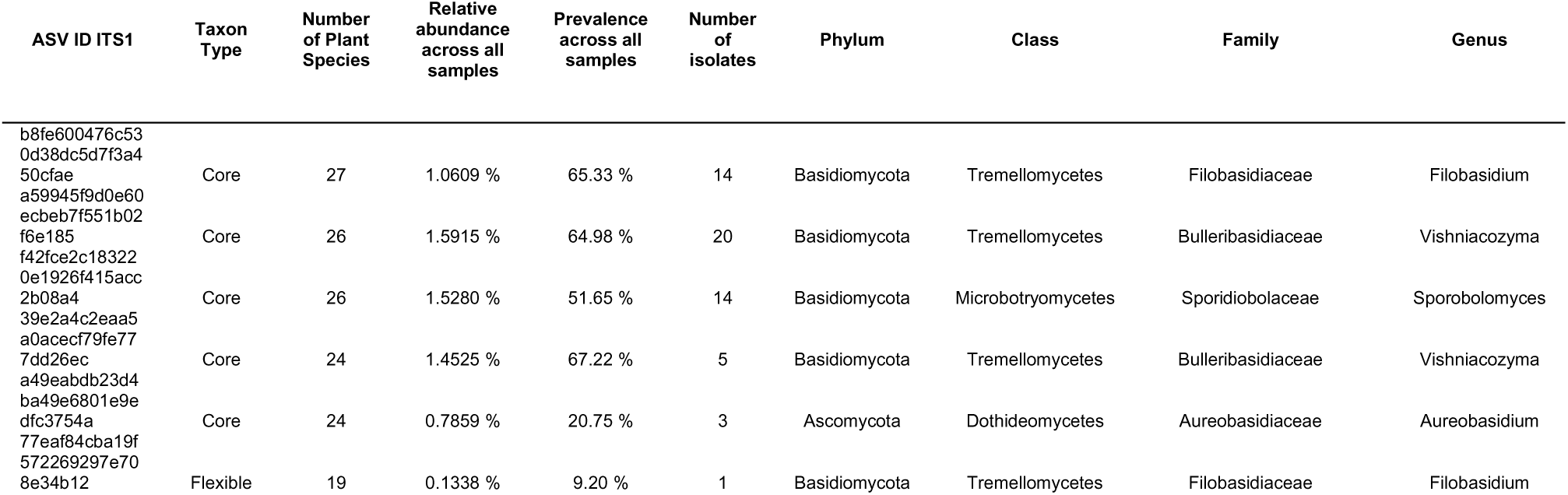
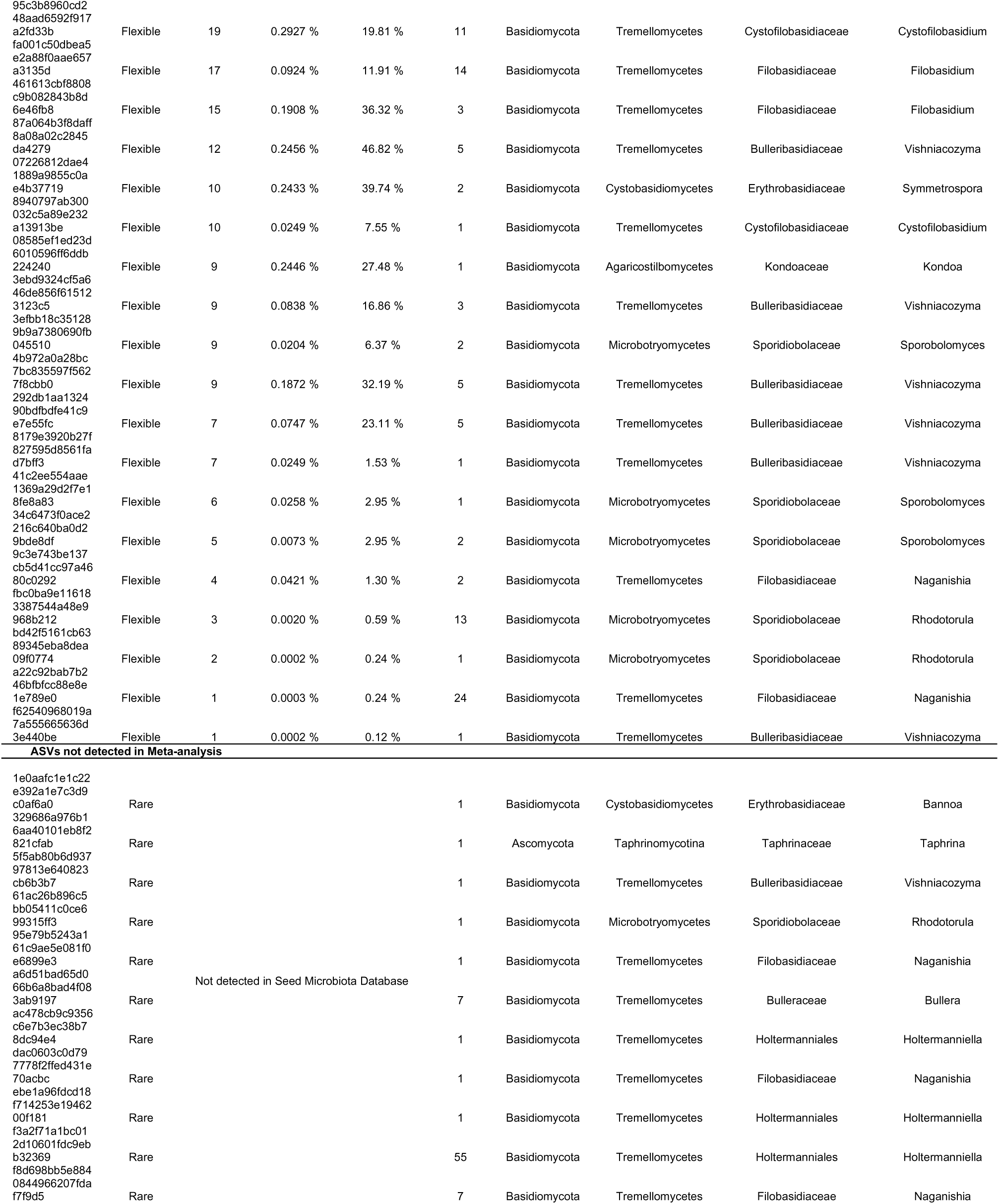
List of yeast ASVs detected or not in Seed Microbiota Database. The 25 detected ASVs in the database collectively represent 8.4% of the relative abundance of the meta-analysis dataset and include 5 core ASVs (out of the six yeast core ASVs) detected in 24 to 27 plant species.

In the seed microbiota meta-analysis, core ASVs were identified based on the arbitrary criterion of a detection in more than 20 plant species (Simonin et al., 2022). Sixteen fungal core ASVs were identified, including six yeast ASVs (3 *Vishniacozyma*, 1 *Filobasidium*, 1 *Sporobolomyces*, 1 *Aureobasidium*). Out of the six core ASVs, five of them were present in our yeast collection (1 Vishniacozyma ASV missing) and we often obtained multiple isolates for these taxa (3 to 20). The other 20 ASVs detected in the meta-analysis were considered as flexible taxa given their lower prevalence and generally low relative abundance i.e. the proportion of these ASVs compared to all others (< 0.3% across all samples). Interestingly, a significant fraction of the isolate diversity (77 isolates, 11 ASVs) had not been previously detected in the Seed Microbiota Database (Table 1) suggesting their rarity or association to specific hosts or environments. Most non-detected ASVs were represented by a single isolate, except for a *Holtermanniella* ASV for which we obtained 55 isolates from different plant species.

## Discussion

Based on our culturomic survey and comparisons with metabarcoding analyses, seed-associated yeasts largely belong to Basidiomycota and more particularly to the Tremellomycetes class. The most frequently isolated yeasts are affiliated with *Holtermanniella*, *Vishniacozyma*, *Filobasidium* and *Sporobolomyces*, which is consistent with previous seed metabarcoding studies (Morales Moreira et al., 2021). This observation is also consistent with the fact that these taxa were recently identified as members of the core seed microbiome (Simonin et al., 2022).

Most of our isolates (n=133) were identified only to the genus level, reflecting the cryptic diversity within fungal communities (Cunningham et al., 2024). While recent studies have updated the taxonomy of Tremellomycetes and continue to describe new species (Gungprakhon et al., 2025; Liu et al., 2025,2015, Wei et al., 2025), further investigations are required to achieve species-level identification of our isolates. Such efforts will not only help fill gaps in the description of seed-associated fungal diversity but also enhance our understanding of microbial dispersion within the plant holobiont. Our strain collection contains multiple isolates sharing the same ITS1 sequences with these core taxa, indicating their high prevalence and abundance across diverse plant hosts and environments. The phylogenetic tree includes 23 sequences from wheat phyllosphere strains from Gouka et al. (2022) study. Among them, 6 strains (F174, F57, F52, F376, F331, F181) are phylogenetically close to seed and seedling isolates within *Sporobolomyces* species, *Vishniacozyma carnescens*, *Holtermaniella takashimae*, *Holtermaniella festucosa*, *Aureobasidium* species. These results suggest that some members of the plant yeast community could persist during plant development and colonize both leaves and seeds. These results encourage further investigation into the potential vertical transmission of yeasts from other aboveground plant compartments to seeds.

However, compared to previous phyllosphere, fruit or flower yeast diversity surveys (Gouka et al., 2022), the yeast from Ascomycota phylum are less frequent in seeds and only represented here by the *Aureobasidium* genus (which is also a seed core taxon) (Simonin et al., 2022) and *Taphrina* sp. 94% isolates from the collection described here belong to Basidiomycota phylum, including red-pigmented isolates. Their coloration is attributed to the production of red, non-diffusible carotenoid pigments, which are abundantly synthesized in strains from *Rhodotorula, Sporobolomyces and Symmetrospora* genera (Frengova & Beshkova, 2009). Carotenoids are thought to protect yeast against oxidative stress, particularly under high UV exposure that they can experience on plant surfaces. This is in line with previous work highlighting that oxidative and hydric stress resistance are crucial for seed microbial colonization (Iacomi-Vasilescu et al., 2008; Pochon et al., 2013). However, determinants conferring yeast seed colonization success remain largely unknown and further work is needed to identify their specific molecular mechanisms and the potential role of morphogenetic adaptations (switch from hyphae to yeast).

In addition to ubiquitous yeast genera, our collection also contains isolates which are specific to one particular plant species. The clearest examples are the case of isolates belonging to *Naganishia* and *Rhodotorula* genera which are specific to *Solanum lycopersicum L.* seeds. This specificity might be due to the wet fruit and seed type of *Solanum lycopersicum L.* that represents a very distinct habitat compared to the other dry seeds included in the survey. The members of *Rhodotorula* and *Naganishia* genera are commonly isolated from different types of environments: water, air, milk etc (Mussagy et al., 2022; Turchetti et al., 2008; Zhao et al., 2019) and not specifically of plant habitats.

Altogether, these results suggest that yeasts are generally well-adapted to the aboveground habitats of plants, but that seeds represent a specific habitat that diverse Basidiomycota yeasts can colonize.

This study presents a previously unexplored yeast diversity on seeds with many genera that had never been observed or isolated in this habitat before. These findings encourage future studies to understand the ecological roles of these yeasts for plant health, especially of the taxa identified as part of the core seed microbiome. The fact that these core taxa are present on numerous plant hosts and environments, suggest a potential co-evolution between the plant and these microbiota members (Risely, 2020). Moreover, some core members and two other genera (*Holtermanniella*, *Vishniacozyma*, *Filobasidium*, *Naganishia*, *Rhodotorula* and *Sporobolomyces*) were isolated from seedlings germinated *in vitro*, which indicates that they could be transmitted between generations from seeds to seedlings. Limited knowledge is available on the persistence and role of yeasts during plant microbiota assembly from seed to seed during two plant generations, but these first results encourage further investigations.

From a more applied point of view, these yeast taxa and especially the ones transmitted to the seedlings are of interest for performing plant microbiota engineering through seed inoculations of single strains or synthetic communities (Arnault et al., 2024; Joubert et al., 2024). Seed yeasts represent an untapped biotechnological resource for seed agriculture that currently mainly rely on bacteria and filamentous fungi (e.g. *Trichoderma*, *Beauvaria*) for biostimulation and biocontrol (Kepler et al., 2017). The relative ease of cultivating and manipulating these yeasts, compared to other fungi, opens numerous promising opportunities for large-scale production of valuable metabolites or field inoculations. Leveraging the deep expertise of the food and beverage industry, yeast-based innovations hold great potential for advancing seed technology, particularly in the areas of crop protection and biofertilizer development.

## Authors contribution

M. Marchi, N. Guschinskaya and M. Simonin contributed to the conception and design, analysis, interpretation of data presented in the publication, as well as to the drafting of the manuscript. S. Aligon, T. Lerenard and J. Le corff harvested seeds from the wild Brassicaceae and isolated microorganisms from them. M. Simonin, M. Briand and C. Marais obtained and analyzed the metabarcoding data. A. Bosc-Bierne, L. Labourgade, M. Marchi and N. Guschinskaya isolated yeast from crops seed. A. Bosc-Bierne and M Marchi obtained molecular typing and M. Marchi carried out the taxonomic affiliation. A. Bosc-Bierne, N. Guschinskaya and A. Rolland contributed to the macroscopic and microscopic characterization of the yeast collection. L Gouka, V. Cordovez, T. Guillemette and P. Simoneau revised the draft manuscript.

## Acknowledgements.

The authors gratefully acknowledge the technical platforms ANAN (Analysis of Nucleic Acids) and IMAC (Cell Imaging) of the SFR QUASAV (Angers, France) for their support and skillful technical assistance, with special thanks to Aurelia Rolland for her valuable advice on image acquisition.

Thanks to the SUCSEED project (ANR-20-PCPA-0009) funded by the ‘Growing and Protecting crops Differently” French Priority Research Program (PPR-PCA), part of the national investment plan operated by the French National Research Agency (ANR) which funded this work.

